# Susceptibility of ferrets, cats, dogs, and different domestic animals to SARS-coronavirus-2

**DOI:** 10.1101/2020.03.30.015347

**Authors:** Jianzhong Shi, Zhiyuan Wen, Gongxun Zhong, Huanliang Yang, Chong Wang, Renqiang Liu, Xijun He, Lei Shuai, Ziruo Sun, Yubo Zhao, Libin Liang, Pengfei Cui, Jinliang Wang, Xianfeng Zhang, Yuntao Guan, Hualan Chen, Zhigao Bu

## Abstract

Severe acute respiratory syndrome coronavirus 2 (SARS-CoV-2) causes the infectious disease COVID-19, which was first reported in Wuhan, China in December, 2019. Despite the tremendous efforts to control the disease, COVID-19 has now spread to over 100 countries and caused a global pandemic. SARS-CoV-2 is thought to have originated in bats; however, the intermediate animal sources of the virus are completely unknown. Here, we investigated the susceptibility of ferrets and animals in close contact with humans to SARS-CoV-2. We found that SARS-CoV-2 replicates poorly in dogs, pigs, chickens, and ducks, but efficiently in ferrets and cats. We found that the virus transmits in cats via respiratory droplets. Our study provides important insights into the animal reservoirs of SARS-CoV-2 and animal management for COVID-19 control.

---

In late December 2019, an unusual pneumonia emerged in humans in Wuhan, China, and rapidly spread internationally, raising global public health concerns. The causative pathogen was identified as a novel coronavirus (*1–16*) that was named Severe Acute Respiratory Syndrome Coronavirus 2 (SARS-CoV-2) on the basis of a phylogenetic analysis of related coronaviruses by the Coronavirus Study Group of the International Committee on Virus Taxonomy (*17*); the disease it causes was subsequently designated COVID-19 by the World Health Organization (WHO). Despite tremendous efforts to control the COVID-19 outbreak, the disease is still spreading. As of March 11, 2020, SARS-CoV-2 infections have been reported in more than 100 countries, and 118,326 human cases have been confirmed, with 4,292 fatalities (*18*). COVID-19 has now been announced as a pandemic by WHO.

Although SARS-CoV-2 shares 96.2% identity at the nucleotide level with the coronavirus RaTG13, which was detected in horseshoe bats (*Rhinolophus* spp) in Yunnan province in 2013 (*3*), it has not previously been detected in humans or other animals. The emerging situation raises many urgent questions. Could the widely disseminated viruses transmit to other animal species, which then become reservoirs of infection? The SARS-CoV-2 infection has a wide clinical spectrum in humans, from mild infection to death, but how does the virus behave in other animals? As efforts are made for vaccine and antiviral drug development, which animal(s) can be used most precisely to model the efficacy of such control measures in humans? To address these questions, we evaluated the susceptibility of different model laboratory animals, as well as companion and domestic animals to SARS-CoV-2.

All experiments with infectious SARS-CoV-2 were performed in the biosafety level 4 and animal biosafety level 4 facilities in the Harbin Veterinary Research Institute (HVRI) of the Chinese Academy of Agricultural Sciences (CAAS), which was approved for such use by the Ministry of Agriculture and Rural Affairs of China. Details of the biosafety and biosecurity measures taken are provided in the supplementary materials (*19*). The protocols for animal study and animal welfare were reviewed and approved by the Committee on the Ethics of Animal Experiments of the HVRI of CAAS.

Ferrets are commonly used as an animal model for respiratory viruses that infected humans (*20–26*). We therefore tested the susceptibility of SARS-CoV-2 in ferrets. Two viruses [SARS-CoV-2/F13/environment/2020/Wuhan, isolated from an environmental sample collected in the Huanan Seafood Market in Wuhan (F13-E), and SARS-CoV-2/CTan/human/2020/Wuhan (CTan-H), isolated from a human patient] were used in this study. Pairs of ferrets were inoculated intranasally with 10^5^ pfu of F13-E or CTan-H, respectively, and euthanized on day 4 post-inoculation (p.i.). The nasal turbinate, soft palate, tonsils, trachea, lung, heart, spleen, kidneys, pancreas, small intestine, brain, and liver from each ferret were collected for viral RNA quantification by qPCR and virus titration in Vero E6 cells. Viral RNA (Fig. 1A, B) and infectious virus were detected in the nasal turbinate, soft palate, and tonsils of all four ferrets inoculated with these two viruses, but was not detected in any other organs tested (Fig. 1C, D). These results indicate that SARS-CoV-2 can replicate in the upper respiratory tract of ferrets, but its replication in other organs is undetectable.

**Figure 1.**
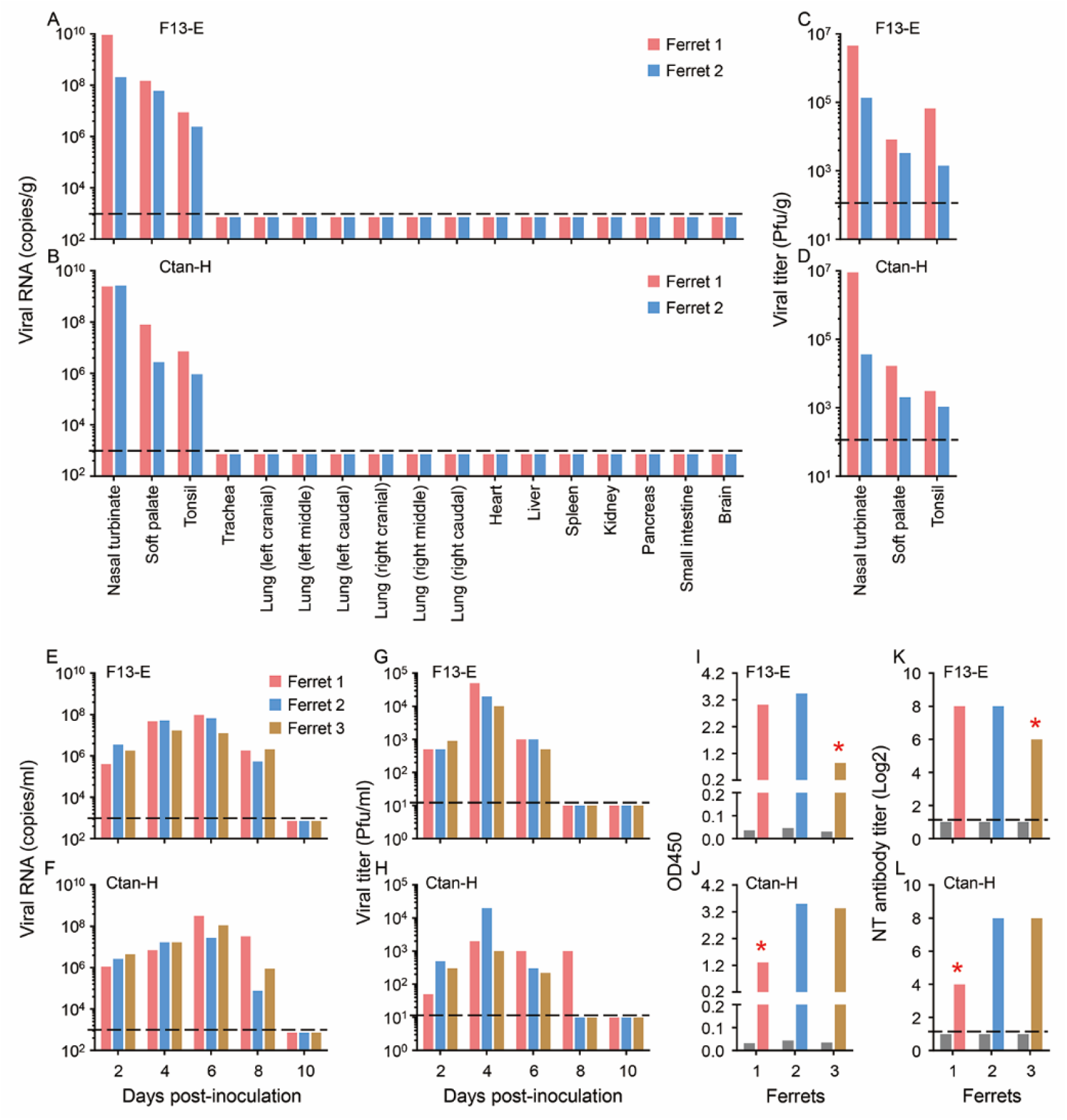
Replication of SARS-CoV-2 viruses in ferrets. Viral RNA in organs or tissues of ferrets inoculated with (**A**) F13-E virus or (**B**) CTan-H virus. Viral titers in organs or tissues of ferrets inoculated with F13-E (**C**) and CTan-H (**D**). Viral RNA in nasal washes of ferrets inoculated with F13-E (**E**) and CTan-H (**F**). Viral titer in nasal washes of ferrets inoculated with F13-E (**G**) and CTan-H (**H**). Antibodies against SARS-CoV-2 tested by an ELISA (**I**, **J**) and neutralization assay (**K, L**) with the sera derived from ferrets inoculated with F13-E (**I, K**) and CTan (**J, L**). Each color bar represents the value from an individual animal. The black bars in the panels **I** to **L** indicate the antibody values of sera collected from each animal before the virus was inoculated. Asterisks indicate animals that were euthanized on day 13 after virus inoculation, the other four animals were euthanized on day 20 p.i. The horizontal dashed lines indicate the lower limit of detection.

To investigate the replication dynamics of these viruses in ferrets, groups of three animals were inoculated intranasally with 10^5^ pfu of F13-E or CTan-H, and then placed in three separate cages within an isolator. Nasal washes and rectal swabs were collected on days 2, 4, 6, 8, and 10 p.i. from the ferrets for viral RNA detection and virus titration. Body temperatures and signs of disease were monitored for two weeks. As shown in Fig. 1, viral RNA was detected in the nasal washes on days 2, 4, 6, and 8 p.i. in all six ferrets inoculated with the two viruses (Fig. 1E, F). Viral RNA was also detected in some of the rectal swabs of the virus-inoculated ferrets although the copy numbers were notably lower than those in the nasal washes of these ferrets (fig. S1A, B). Infectious virus was detected from the nasal washes of all ferrets (Fig. 1G, H), but not from the rectal swabs of any ferrets (fig. S1C, D).

One ferret from each virus-inoculated group developed fever and loss of appetite on days 10 and 12 p.i., respectively. To investigate whether these symptoms were caused by virus replication in the lower respiratory tract, we euthanized the two ferrets on day 13 p.i., and collected their organs for viral RNA detection. However, viral RNA was not detected in any other tissues or organs of either ferret, except for a low copy number (10^5.4^ copies/g) in the turbinate of the ferret inoculated with CTan-H (fig S2). Pathological studies revealed severe lymphoplasmacytic perivasculitis and vasculitis, increased numbers of type II pneumocytes, macrophages, and neutrophils in the alveolar septa and alveolar lumen, and mild peribronchitis in the lungs of the two ferrets euthanized on day 13 p.i. (fig.S3). Antibodies against SARS-CoV-2 were detected in all of the ferrets by an ELISA and a neutralization assay, although the antibody titers of the two ferrets that were euthanized on day 13 p.i. were notably lower than those of the ferrets euthanized on day 20 p.i. (Fig. 1I-L).

To further investigate whether SARS-CoV-2 replicates in the lungs of ferrets, we intratracheally inoculated eight ferrets with 10^5^ pfu of CTan-H, and euthanized two animals each on days 2, 4, 8, and 14 p.i. to look for viral RNA in the tissues and organs. Viral RNA was only detected in the nasal turbinate and soft palate of one of the two ferrets that were euthanized on days 2 and 4 p.i.; in the soft palate of one ferret and in the nasal turbinate, soft palate, tonsil, and trachea of the other ferret that were euthanized on day 8 p.i.; and was not detected in either of the two ferrets that were euthanized on day 14 p.i. (fig. S4). These results indicate that SARS-CoV-2 can replicate in the upper respiratory tract of ferrets for up to eight days, without causing severe disease or death.

Cats and dogs are in close contact with humans, and therefore it is important to understand their susceptibility to SARS-CoV-2 for COVID-19 control. We first investigated the replication of SARS-CoV-2 in cats. Five 8-month-old outbred domestic cats (referred to here as “subadult cats”) were intranasally inoculated with 10^5^ pfu of CTan-H. Two of the subadult cats were scheduled to be euthanized on day 6 p.i. to evaluate viral replication in their organs. Three subadult cats were placed in separate cages within an isolator. To monitor respiratory droplet transmission, an uninfected cat was placed in a cage adjacent to each of the infected cats. It was difficult to perform regular nasal wash collection on the subadult cats because they were aggressive. To avoid possible injury, we only collected feces from these cats and checked for viral RNA in their organs after euthanasia.

From the two subadult cats that were euthanized on day 6 p.i. with CTan-H, viral RNA was detected in the nasal turbinates, soft palates, and tonsils of both animals, in the trachea of one animal, and in the small intestine of the other; however, viral RNA was not detected in any of the lung samples from either of these animals (Fig. 2A). Infectious virus was detected in the viral RNA-positive nasal turbinates, soft palates, tonsils, and trachea of these cats, but was not recovered from the viral RNA-positive small intestine (Fig. 2B).

**Figure 2.**
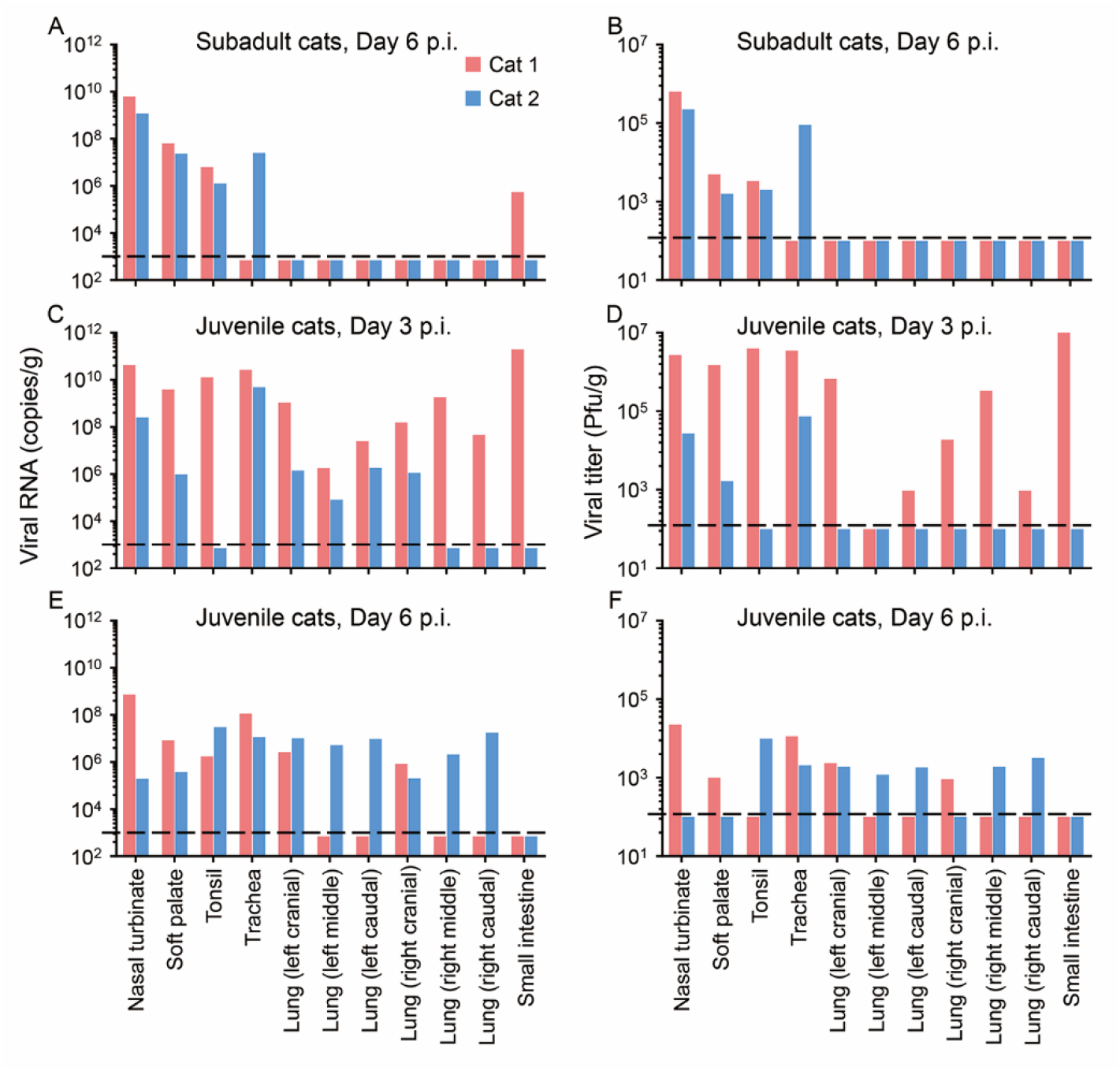
Replication of SARS-CoV-2 in cats. (**A**) Viral RNA and (**B**) viral titers detected in subadult cats that were euthanized on day 6 post-inoculation (p.i.) with CTan-H virus. (**C**) Viral RNA and (**D**) viral titers detected in juvenile cats that were euthanized on day 3 p.i. with CTan-H virus. (**E**) Viral RNA and (**F**) viral titers detected in juvenile cats that were euthanized on day 6 p.i. with CTan-H virus. Each color bar represents the value from an individual animal. The horizontal dashed lines indicate the lower limit of detection.

In the transmission study, viral RNA was detected in the feces of two virus-inoculated subadult cats on day 3 p.i., and in all three virus-inoculated subadult cats on day 5 p.i. (Fig. 3A). Viral RNA was detected in the feces of one exposed cat on day 3 p.i. (Fig. 3A). The subadult cats with viral RNA-positive feces were euthanized on day 11 p.i., and viral RNA was detected in the soft palate and tonsils of the virus-inoculated animal and in the nasal turbinate, soft palate, tonsils, and trachea of the exposed animal (Fig. 3B), indicating that respiratory droplet transmission had occurred in this pair of cats. We euthanized the other pairs of animals on day 12 p.i., and viral RNA was detected in the tonsils of one virus-inoculated subadult cat, in the nasal turbinate, soft palate, tonsils, and trachea of the other virus-inoculated subadult cat, but was not detected in any organs or tissues of the two exposed subadult cats (Fig. 3B). Antibodies against SARS-CoV-2 were detected in all three virus-inoculated subadult cats and one exposed cat by use of an ELISA and neutralization assay (Fig. 3C, D).

**Figure 3.**
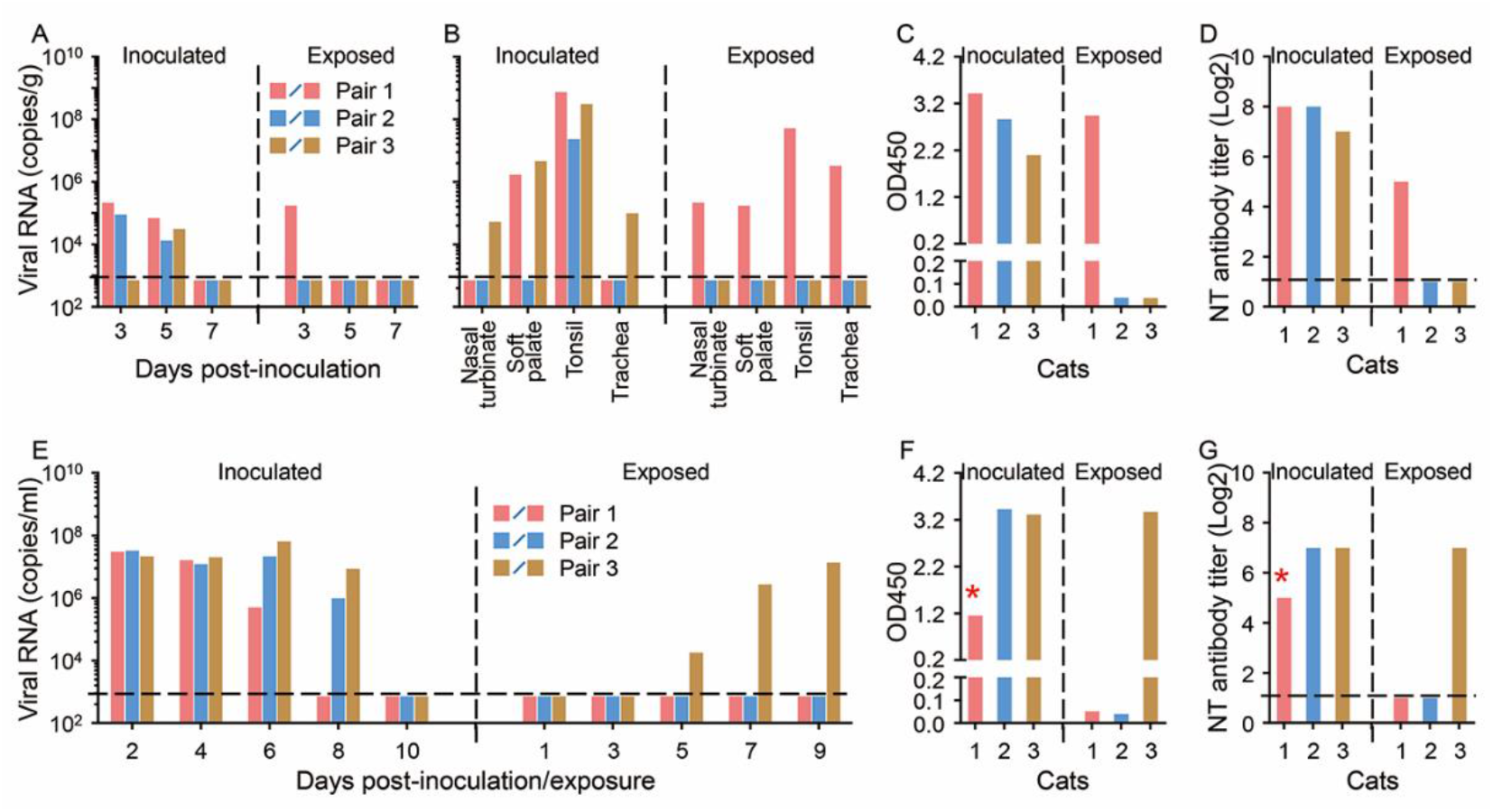
Transmission of SARS-CoV-2 in cats. (**A**) Viral RNA in the feces of subadult cats that were inoculated with or exposed to CTan-H at the indicated timepoints. (**B**) Viral RNA in tissues or organs of subadult cats that were inoculated with or exposed to CTan-H, the pair one cats (red bars) were euthanized on day 11 p.i. and the other two pairs were euthanized on day 12 p.i. Antibodies against SARS-CoV-2 of these euthanized cats were detected by using an ELISA (**C**) and neutralization assay (**D**). (**E**) Viral RNA in nasal washes of juvenile cats that were inoculated with or exposed to CTan-H. Sera of the juvenile cats were collected on day 20 p.i., and their antibodies against SARS-CoV-2 were detected by using an ELISA (**F**) and neutralization assay (**G**). One virus inoculated cat died on day 13 p.i. and the antibody values of this cat (indicated by asterisks) were detected from the sera collected on day 10 p.i. Each color bar represents the value from an individual animal. The horizontal dashed lines indicate the lower limit of detection.

We replicated the replication and transmission studies in juvenile cats (aged 70–100 days) (Fig. 2C-F and Fig. 3E-G). Histopathologic studies performed on samples from the virus-inoculated juvenile cats that died or euthanized on day 3 p.i. revealed massive lesions in the nasal and tracheal mucosa epitheliums, and lungs of both cats (fig. S5). These results indicate that SARS-CoV-2 can replicate efficiently in cats, with younger cats being more permissive and, perhaps more importantly, the virus can transmit between cats via respiratory droplets.

We next investigated the replication and transmission of SARS-CoV-2 in dogs. Five 3-month-old beagles were intranasally inoculated with 10^5^ pfu of CTan-H, and housed with two uninoculated beagles in a room. Oropharyngeal and rectal swabs from each beagle were collected on days 2, 4, 6, 8, 10, 12, and 14 p.i., for viral RNA detection and virus titration in Vero E6 cells. Viral RNA was detected in the rectal swabs of two virus-inoculated dogs on day 2 p.i., and in the rectal swab of one dog on day 6 p.i. (Table 1). One dog that was viral RNA positive by its rectal swab on day 2 p.i. was euthanized on day 4 p.i., but viral RNA was not detected in any organs or tissues collected from this dog (fig. S6). Infectious virus was not detected in any swabs collected from these dogs. Sera were collected from all of the dogs on day 14 p.i. for antibody detection by use of an ELISA. Two virus-inoculated dogs seroconverted; the other two virus-inoculated dogs and the two contact dogs were all seronegative for SARS-CoV-2 according to the ELISA (Table 1, fig S7). These results indicate that dogs have low susceptibility to SARS-CoV-2.

**Table 1.** Susceptibility of dogs, pigs, chickens, and ducks to SARS-CoV-2^a^.

| Animal | Treatment | Viral RNA detection in animals inoculated with SARS-CoV-2 isolate CTan-H; No. of positive/total (copies, log<sub>10</sub>) |  |  |  |  |  |  |  | Seroconversion: |
| --- | --- | --- | --- | --- | --- | --- | --- | --- | --- | --- |
|  |  | Oropharyngeal swab |  |  |  | Rectal swab |  |  |  | No. of positive |
|  |  | Day 2 p.i. | Day 4 p.i. | Day 6 p.i. | Other timepoints | Day 2 p.i. | Day 4 p.i. | Day 6 p.i. | Other timepoints | /Total<sup>c</sup> |
| Dog<sup>b</sup> | Inoculated | 0/5 | 0/5 | 0/4 | 0/4 | 2/5 (6.5, 5.4) | 0/5 | 1/4 (4.2) | 0/4 | 2/4 |
|  | Contact | 0/2 | 0/2 | 0/2 | 0/2 | 0/2 | 0/2 | 0/2 | 0/2 | 0/2 |
| Pig | Inoculated | 0/5 | 0/5 | 0/5 | 0/5 | 0/5 | 0/5 | 0/5 | 0/5 | 0/5 |
|  | Contact | 0/3 | 0/3 | 0/3 | 0/3 | 0/3 | 0/3 | 0/3 | 0/3 | 0/3 |
| Chicken | Inoculated | 0/5 | 0/5 | 0/5 | 0/5 | 0/5 | 0/5 | 0/5 | 0/5 | 0/5 |
|  | Contact | 0/3 | 0/3 | 0/3 | 0/3 | 0/3 | 0/3 | 0/3 | 0/3 | 0/3 |
| Duck | Inoculated | 0/5 | 0/5 | 0/5 | 0/5 | 0/5 | 0/5 | 0/5 | 0/5 | 0/5 |
|  | Contact | 0/3 | 0/3 | 0/3 | 0/3 | 0/3 | 0/3 | 0/3 | 0/3 | 0/3 |

- Animals were intranasally inoculated with 10<sup>5</sup> pfu (dogs and pigs) or 10<sup>4.5</sup> pfu (chickens and ducks) of the CTan-H virus, and two (beagles) or three (pigs, chickens, and ducks) uninfected animals were housed in the same room with their infected counterparts to monitor the transmission of the CTan-H virus. Oropharyngeal and rectal swabs from all animals were collected on the indicated days post inoculation (p.i.) for viral RNA detection. Other timepoints include days 8, 10, 12, and 14 p.i.. - One virus-inoculated beagle was euthanized on day 4 p.i., but viral RNA was not detected in any of its collected organs, which included lung, trachea, nasal turbinate, soft palate, brain, heart, submaxillary lymph nodes, tonsil, kidneys, spleen, liver, pancreas, and small intestine (data not shown). - Sera were collected from all animals on day 14 p.i., and antibodies against SARS-Cov-2 were detected by using a Double Antigen Sandwich ELISA kit (ProtTech, Luoyang, China).

We also investigated the susceptibility of pigs, chickens, and ducks to SARS-CoV-2 by using the same strategy as that used to assess dogs; however, viral RNA was not detected in any swabs collected from these virus-inoculated animals or from naïve contact animals (Table 1), and all of the animals were seronegative for SARS-CoV-2 when tested by using the ELISA with sera collected on day 14 p.i. (Table 1). These results indicate that pigs, chickens, and ducks are not susceptible to SARS-CoV-2.

In summary, we found that ferrets and cats are highly susceptible to SARS-CoV-2, dogs have low susceptibility, and livestock including pigs, chickens, and ducks are not susceptible to the virus.

Ferrets have frequently been used as an animal model for the study of human respiratory viruses (*20–26*). Unlike influenza viruses and other human SARS-coronavirus, which replicate in both the upper and lower respiratory tract of ferrets (*20, 22–24, 26, 27*), we found SARS-CoV-2 only replicates in the nasal turbinate, soft palate, and tonsils of ferrets. It may also replicate in the digestive tract, as viral RNA was detected in the rectal swabs of the virus-infected ferrets, but virus was not detected in lung lobes, even after the ferrets were intratracheally inoculated with the virus.

Several studies have reported that SARS-CoV-2 uses angiotensin-converting enzyme 2 (ACE2) as its receptor to enter cells (*3, 28–30*). ACE2 is mainly expressed in type II pneumocytes and serous epithelial cells of tracheo-bronchial submucosal glands in ferrets (*25*); therefore, the underlying mechanism that prevents the replication of SARS-CoV-2 in the lower respiratory tract of ferrets remains to be investigated. The fact that SARS-CoV-2 replicates efficiently in the upper respiratory tract of ferrets makes them a candidate animal model for evaluating antiviral drugs or vaccine candidates against COVID-19.

The cats we used in this study were outbred, and were highly susceptible to SARS-CoV-2, which replicated efficiently and transmitted to naïve cats. Surveillance for SARS-CoV-2 in cats should be considered as an adjunct to elimination of of COVID-19 in humans.

## Acknowledgments

We thank S. Watson for editing the manuscript. This work was supported by the National Key R&D Program of China (2018YFC1200601). We thank professors Wenjie Tan and Guizhen Wu from China CDC for providing the SARS-CoV-2 strains. This work is licensed under a Creative Commons Attribution 4.0 International (CC BY 4.0) license, which permits unrestricted use, distribution, and reproduction in any medium, provided the original work is properly cited. To view a copy of this license, visit https://creativecommons.org/licenses/by/4.0/. This license does not apply to figures/photos/artwork or other content included in the article that is credited to a third party; obtain authorization from the rights holder before using such material.

## Author contributions

J.S., Z.W., G.Z., H.Y., C.W., R.L., X.H., L.S., Z.S., Y.Z., L.L., P.C., J.W., X.Z., and Y.G. performed experiments; J.S., Z.W., G.Z., H.Y., C.W., Z.B., and H.C. analyzed data; Z.B. and H.C. designed the study and wrote the manuscript.

## Competing interests

None of the authors has any competing interests.

## Data and materials availability

all data is available in the manuscript or the supplementary materials.

## Supporting information

### Text

Histopathologic and immunohistochemical studies were performed on samples from the virus-inoculated juvenile cats that died or euthanized on day 3 p.i. The nasal respiratory mucosa epithelium exhibited an abnormal arrangement with loss of cilia accompanied by lymphocyte infiltration into the lamina propria (fig. S5A). In the tonsils, the surface of the epithelium was covered with abundant neutrophils and cellular debris, and the epithelial cells showed varying degrees of degeneration and necrosis (fig. S5B). The tracheal mucosa was covered with cellular debris and mucous material, altered polarity, degeneration, and necrosis was observed in the epithelial cells (fig. S5C), and epithelial necrosis and lymphocyte infiltration were observed in the submucosal glands (fig. S5D). Inflammatory cell (including neutrophils, mononuclear cells, lymphocytes) aggregation and fibrin formation in the lung vasculature were seen (fig. S5E), infiltration of a large number of macrophages and lymphocytes into the alveolar spaces and interalveolar septa (fig. S5F), and intra-alveolar edema and congestion in the interalveolar septa were commonly observed (fig. S5G). In the small intestine, diffuse degeneration of mucosal epithelial cells was observed, some of which showed necrosis, and moderate lymphocytic infiltration was observed in the mucosal lamina propria (fig. S5H). Large amounts of viral antigen were detected in the epithelium of the nasal respiratory mucosa (fig. S5I) and in the epithelial cells of the tonsils; tonsillar surficial cellular debris was evident (fig. S5J). Although viral antigen was hardly detected in the tracheal epithelial cells, it was highly apparent in the serous cells of the tracheo-bronchial submucosal glands (fig. S5K). Numerous epithelial cells of the small intestine also showed strong positive staining for viral antigen (fig. S5L).

### MATERIALS AND METHODS

#### Facility, Ethics, and Biosafety statement

All experiments with infectious SARS-CoV-2 were performed in the biosafety level 4 and animal biosafety level 4 facilities in the Harbin Veterinary Research Institute (HVRI) of the Chinese Academy of Agricultural Sciences (CAAS), which is approved for such use by the Ministry of Agriculture and Rural Affairs of China. The animal studies were carried out in strict accordance with the recommendations in the Guide for the Care and Use of Laboratory Animals of the Ministry of Science and Technology of the People’s Republic of China. The protocols were approved by the Committee on the Ethics of Animal Experiments of the HVRI of CAAS.

All experiments were conducted within the biosafety level 4 (P4) facilities in the Harbin Veterinary Research Institute (HVRI) of the Chinese Academy of Agricultural Sciences (CAAS), which was completed in 2015 and accredited by the China National Accreditation Service for Conformity Assessment in 2018. Handling of SARS-CoV-2 was approved by the Ministry of Agriculture and Rural Affairs of China on January 22 and by the National Health Commission of China on January 31. All activities inside the P4 labs are monitored by trained guards via video cameras. Only authorized personnel that have received appropriate training can access the P4 facility. Experienced personnel work in pairs in the facilities. Our staff wear powered air-purifying respirators that filter the air, and disposable coveralls when they culture the virus, and wear full body, air-supplied, positive pressure suits when they perform the animal study; they were disinfected before they leave the room and then shower on exiting the facility. The facility is secured by appropriate procedures approved by the HRVI institutional biosafety officers. All facilities, procedures, training records, safety drills, and inventory records are subject to periodic inspections and ongoing oversight by the institutional biosafety officers who consult frequently with the facility managers. The research program, procedures, occupational health plan, security and facilities are reviewed annually by a Ministry of Agriculture and Rural Affairs official.

#### Cells

Vero-E6 cells were maintained in Dulbecco’s modified Eagle’s medium (DMEM) containing 10% fetal bovine serum (FBS) and antibiotics and incubated at 37°C with 5% CO_2_.

#### Virus

SARS-CoV-2/CTan/human/2020/Wuhan (CTan-H) was isolated from a human patient and SARS-CoV-2/F13/environment/2020/Wuhan (F13-E, formally designated as SARS-CoV-2/F13/environment/2020/Wuhan, GISAID access no. EPI_ISL_408515) virus was isolated from an environmental sample collected in Huanan seafood market, Wuhan, and were kindly provided by professor Guizhen Wu from the National Institute for Viral Disease Control and Prevention, Chinese Center for Disease Control and Prevention. Viral stocks were prepared in Vero E6 cells with DMEM containing 2% FBS, 5 ug/ml TPCK-trypsin, and 30 mmol/L MgCl_2_. Viruses were harvested and the titers were determined by means of plaque assays in Vero E6 cells.

#### qPCR

To quantitate the viral RNA copies in the samples collected from the animals, viral RNA was extracted by using a QIAamp vRNA Minikit (Qiagen, Hilden, Germany). Reverse transcription was performed by using the HiScript^®^ II Q RT SuperMix for qPCR (Vazyme, Nanjing, China). qPCR was conducted by using the Applied Biosystems^®^ QuantStudio^®^ 5 Real-Time PCR System (Thermo Fisher Scientific, Waltham, MA, USA) with Premix Ex Taq™ (Probe qPCR), Bulk (TaRaKa, Dalian, China). The N gene-specific primers (forward, 5’-GGGGAACTTCTCCTGCTAGAAT-3’; reverse, 5’-CAGACATTTTGCTCTCAAGCTG-3) and probe (5’-FAM-TTGCTGCTGCTTGACAGATT-TAMRA-3’) were utilized according to the information provided by the National Institute for Viral Disease Control and Prevention, China (http://nmdc.cn/nCoV). The amount of vRNA for the target SARS-CoV-2 N gene was normalized to the standard curve obtained by using a plasmid (pBluescriptⅡSK-N, 4,221 bp) containing the full-length cDNA of the SARS-CoV-2 N gene.

#### ELISA

Antibodies against SARS-CoV-2 were detected by using a Double Antigen Sandwich ELISA Kit (ProtTech, Luoyang, China) according to the manufacturer’s instructions. Briefly, 100 μL of sera were added to an antigen-precoated microtiter plate and incubated at 37 °C for 30 min. Plates were then washed 5 times with PBST, and incubated with HRP-conjugated antigen at 37 °C for 30 min. Plates were washed 5 times with PBST, and 100 μL of substrate solution was added to enable colorimetric analysis. The reaction was stopped by adding 50 μL of stop buffer, and optical density (OD) was measured at 450 nm.

#### Ferret Study

Three-to four-month-old female ferrets (Wuxi Cay Ferret Farm, Wuxi, China) were used in this study. Groups of five ferrets were anesthetized with Zoletil 50 (Virbac, Carros, France) and inoculated intranasally with 10^5^ PFU of SARS-CoV-2 CTan-H or F13-E in a volume of 1 mL and each ferret was housed in a separate cage inside an isolator. Two ferrets from each group were euthanized on day 4 post-inoculation (p.i.) and their organs and tissues, including lungs, tracheas, nasal turbinates, soft palates, brains, hearts, submaxillary lymph nodes, tonsils, kidneys, spleens, livers, pancreas, and small intestines, were collected for viral RNA and virus detection. Nasal washes and rectal swabs were collected from the other three ferrets on days 2, 4, 6, 8, and 10 p.i. for viral RNA and virus detection. Body weights and body temperature were monitored every other day. The ferrets were scheduled to be euthanized on day 20 p.i. to assess them for antibodies against SARS-CoV-2 by using the Double Antigen Sandwich ELISA kit (ProtTech, Luoyang, China) and by use of a neutralization assay.

Eight ferrets were also inoculated intratracheally with 10^5^ PFU of CTan-H. On days 2, 4, 8, and 14 p.i., two ferrets were each euthanized, and their organs were collected for virusdetection.

#### Cat study

Ten outbred domestic juvenile cats (aged between 70 days and three months) and eight outbred subadult cats (aged 6–9 months) obtained from the National Engineering Research Center of Veterinary Biologics CORP. (Harbin, China) were used in this study. To evaluate the replication of SARS-CoV-2, four juvenile cats and two subadult cats were intranasally inoculated with 10^5^ PFU of CTan-H. Two juvenile cats were euthanized on day 3 p.i.; two other juvenile cats and the two subadult cats were euthanized on day 6 p.i. Organs and tissues were collected for viral RNA detection, virus titration in Vero E6 cells, and histological studies.

To investigate the transmissibility of SARS-CoV-2 in cats, six cats were inoculated intranasally with 10^5^ PFU of CTan-H and each animal was placed in a separate cage within an isolator. A similar-aged naive cat was then placed in each cage adjacent to the one that held the virus-inoculated cat. Feces were collected on days 3, 5, 7, and 9 p.i. from the big cats, and nasal washes were collected from the juvenile cats for viral RNA detection or virus titration. One pair and two pairs of the subadult cats were euthanized on days 11 and 12 p.i., respectively, and their organs, including lungs, tracheas, nasal turbinates, soft palates, brains, hearts, submaxillary lymph nodes, tonsils, kidneys, spleens, livers, pancreas, and small intestines, were collected for viral RNA detection and virus titration. Sera were collected and antibodies against SARS-CoV-2 were detected by using the Double Antigen Sandwich ELISA kit (ProtTech, Luoyang, China) and a neutralization assay.

#### Dog, pig, chickens, and duck study

To access the susceptibility of dogs, pigs, chickens, and ducks to SARS-CoV-2, three-month-old beagles (Kangping Institute, Shenyang, China), 40-day-old specific-pathogen-free (SPF) Landrace and Large White pigs (HVRI, Harbin, China), 4-week-old specific-pathogen-free (SPF) White Leghorn chickens (HVRI, Harbin, China), and 4-week-old SPF ducks (Shaoxin ducks, a local bred) (HVRI, Harbin, China) were used in this study.

Animals were intranasally inoculated with 10^5^ PFU (dogs and pigs) or 10^4.5^ PFU (chickens and ducks) of CTan-H, and two (dogs) or three (pigs, chickens, ducks) uninfected animals were housed in the same room with their infected counterparts to monitor the transmission of CTan-H. Oropharyngeal and rectal swabs from all animals were collected every other day for viral RNA detection. One beagle was viral RNA positive by its rectal swab on day 2 p.i. and was euthanized on day 4 p.i.; its organs, including nasal turbinates, soft palates, tonsils, tracheas, lungs, brains, hearts, kidneys, spleens, livers, pancreas, and small intestines, were collected for viral RNA detection by qPCR. Sera were collected from all remaining animals on day 14 p.i., and antibodies against SARS-CoV-2 were detected by using a Double Antigen Sandwich ELISA kit (ProtTech, Luoyang, China).

#### Plaque reduction neutralization test

SARS-CoV-2 CTan strain (100 PFU) was incubated with two-fold serial dilutions of sera for 1 h at 37°C. A plaque assay was then performed in Vero E6 cells with the neutralization mixtures. Neutralizing antibody titers were calculated as the maximum serum dilution yielding a 50% reduction in the number of plaques relative to control serum prepared from uninfected animals.

#### Histological study

Tissues of animals were fixed in 10% neutral-buffered formalin, embedded in paraffin, and cut into 4-μm sections. The sections were stained with hematoxylin-eosin (H&E) or used in immunohistochemical (IHC) assays. The sections used for immunohistochemistry were dewaxed in xylene and hydrated through a series of descending concentrations of alcohol to water. For viral antigen retrieval, sections were immersed in citric acid/sodium citrate solution at 121 °C for 15 minutes. After cooling, the sections were treated with 3% hydrogen peroxide for 30 minutes to remove endogenous peroxidase activity and blocked with 8% skim milk to reduce nonspecific binding. After three 5-minute washes in TBS, the sections were incubated with rabbit anti-SARS-CoV-2 nucleoprotein monoclonal antibody (1:500; Frdbio. Wuhan, China) in 8% skim milk at 4 °C overnight. The sections were then washed again with TBS and incubated with anti-rabbit IgG (whole molecule)-HRP (1:600, Sigma-Aldrich) at room temperature for 60 minutes. The immunostaining was visualized with DAB and counterstained with hematoxylin.

---

**Figure S1.**
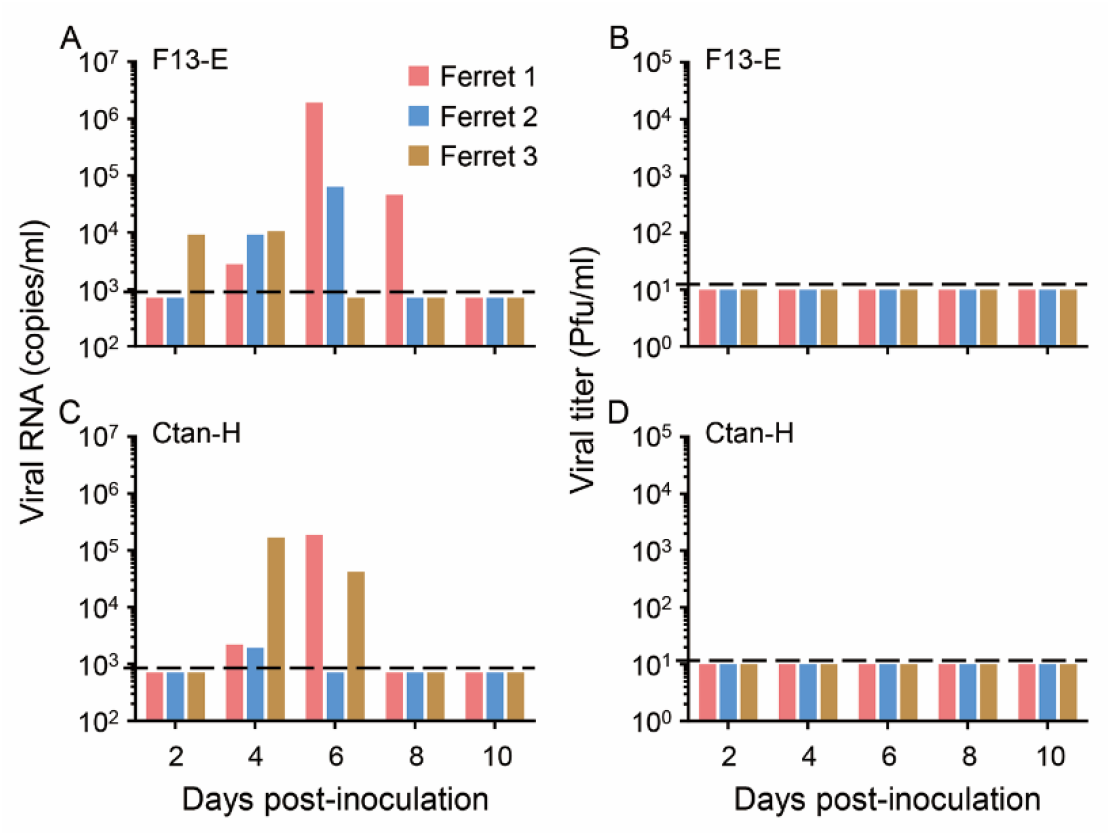
Replication of SARS-CoV-2 in digestive tract of ferrets. Viral RNA in rectal swabs of ferrets inoculated with F13-E (**A**) and CTan-H (**B**). Viral titer in rectal swabs of ferrets inoculated with F13-E (**C**) and CTan-H (**D**).

**Figure S2.**
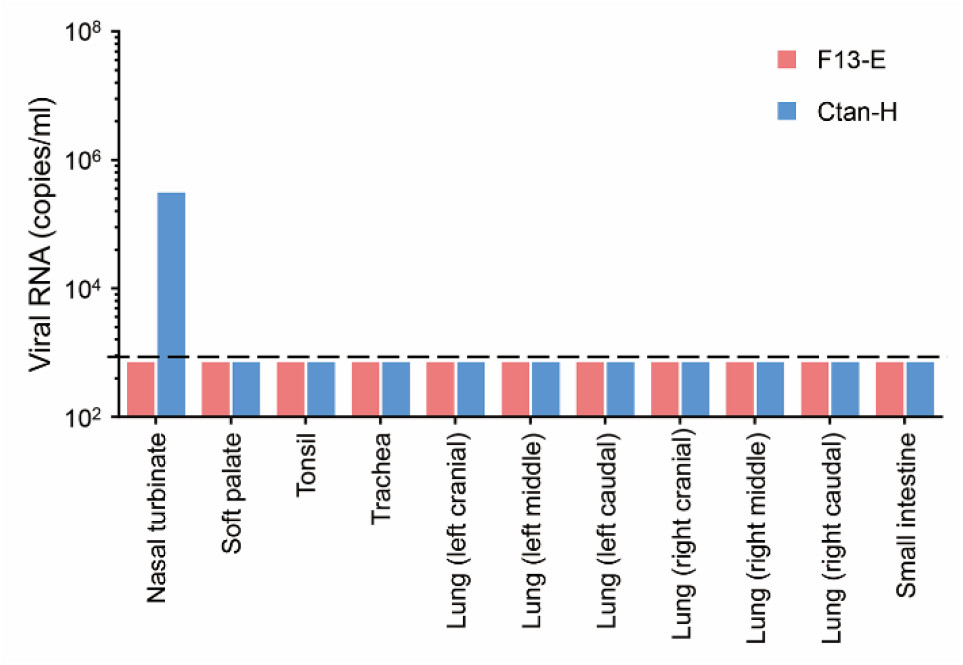
Replication of SARS-CoV-2 in ferrets. Ferrets inoculated with F13-E virus or CTan-H virus were euthanized on day 13 p.i. and their organs and tissues were collected for viral RNA detection.

**Figure S3.**
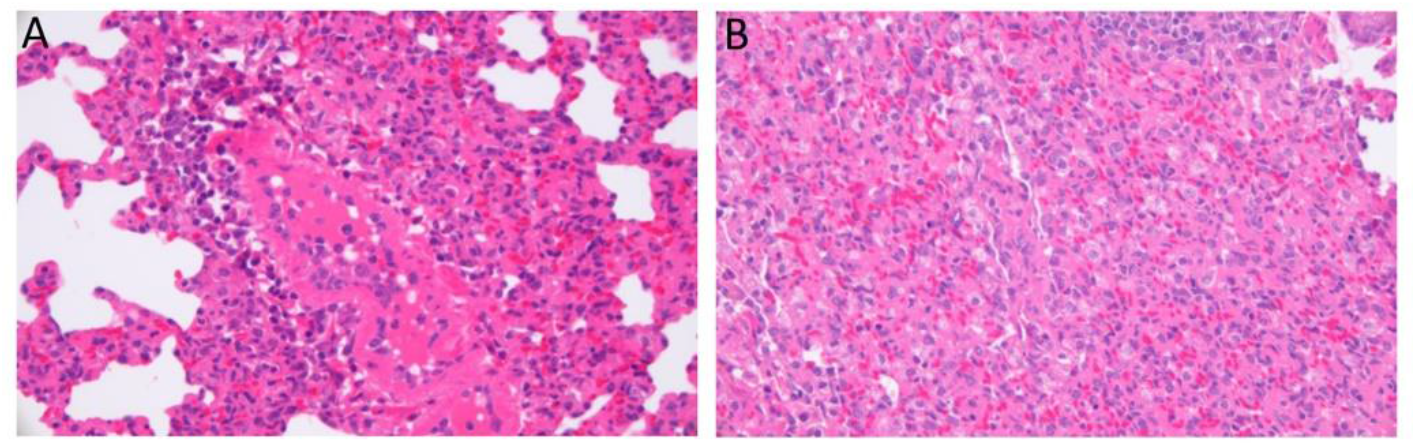
Histological study of a lung sample from a ferret infected with SARS-CoV-2 CTan-H. **(A)** Severe lymphoplasmacytic perivasculitis and vasculitis. **(B)** Increased numbers of Type II pneumocytes, macrophages, and neutrophils in the alveolar septa and alveolar lumen. Original magnifications: ×400

**Figure S4.**
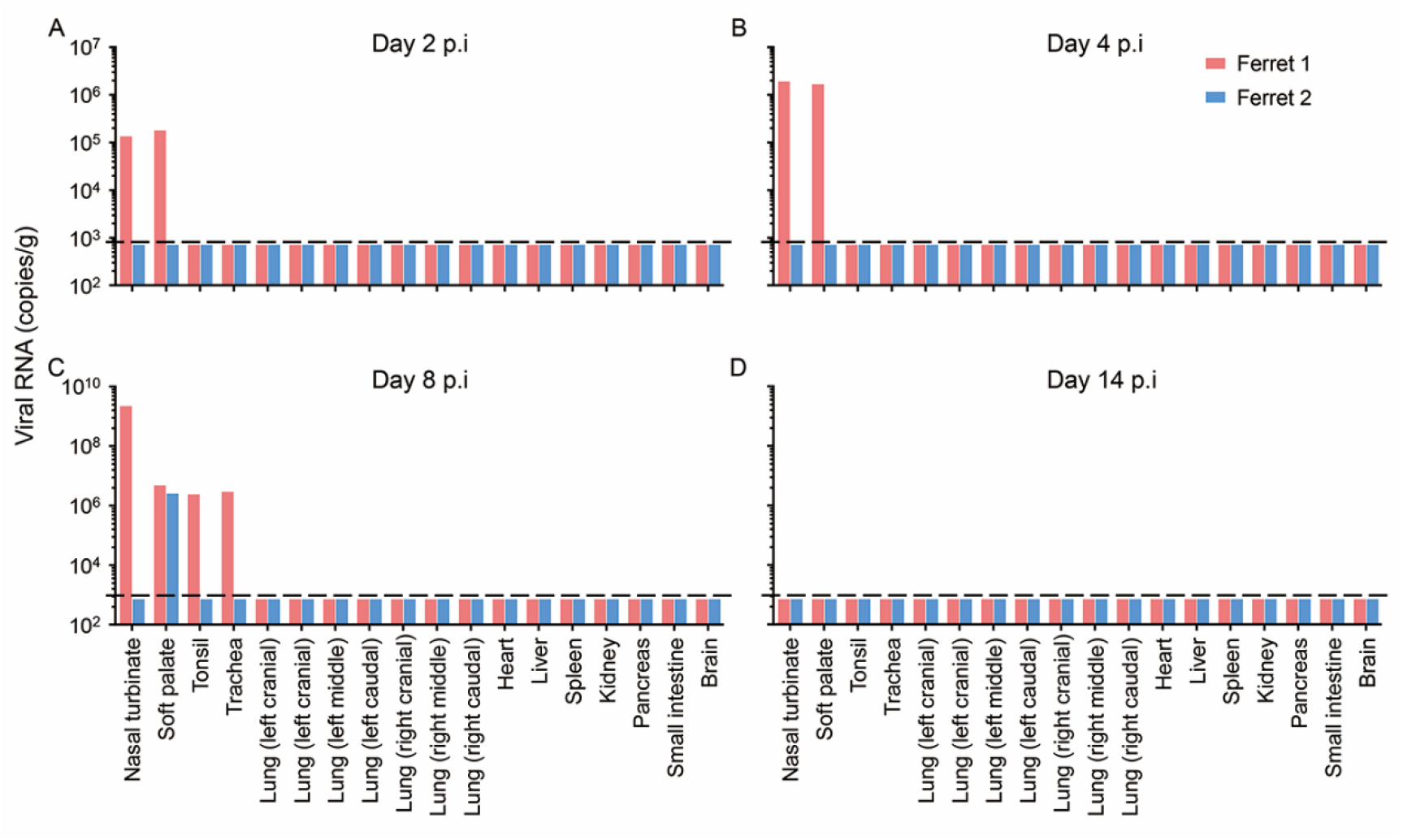
Viral RNA copies in the organs of ferrets intratracheally inoculated with CTan-H. Eight ferrets were inoculated intratracheally with 10^5^ PFU of CTan-H. On days 2, 4, 8, and 14 p.i., two ferrets were each euthanized, and their organs were collected for virus detection by qPCR.

**Figure S5.**
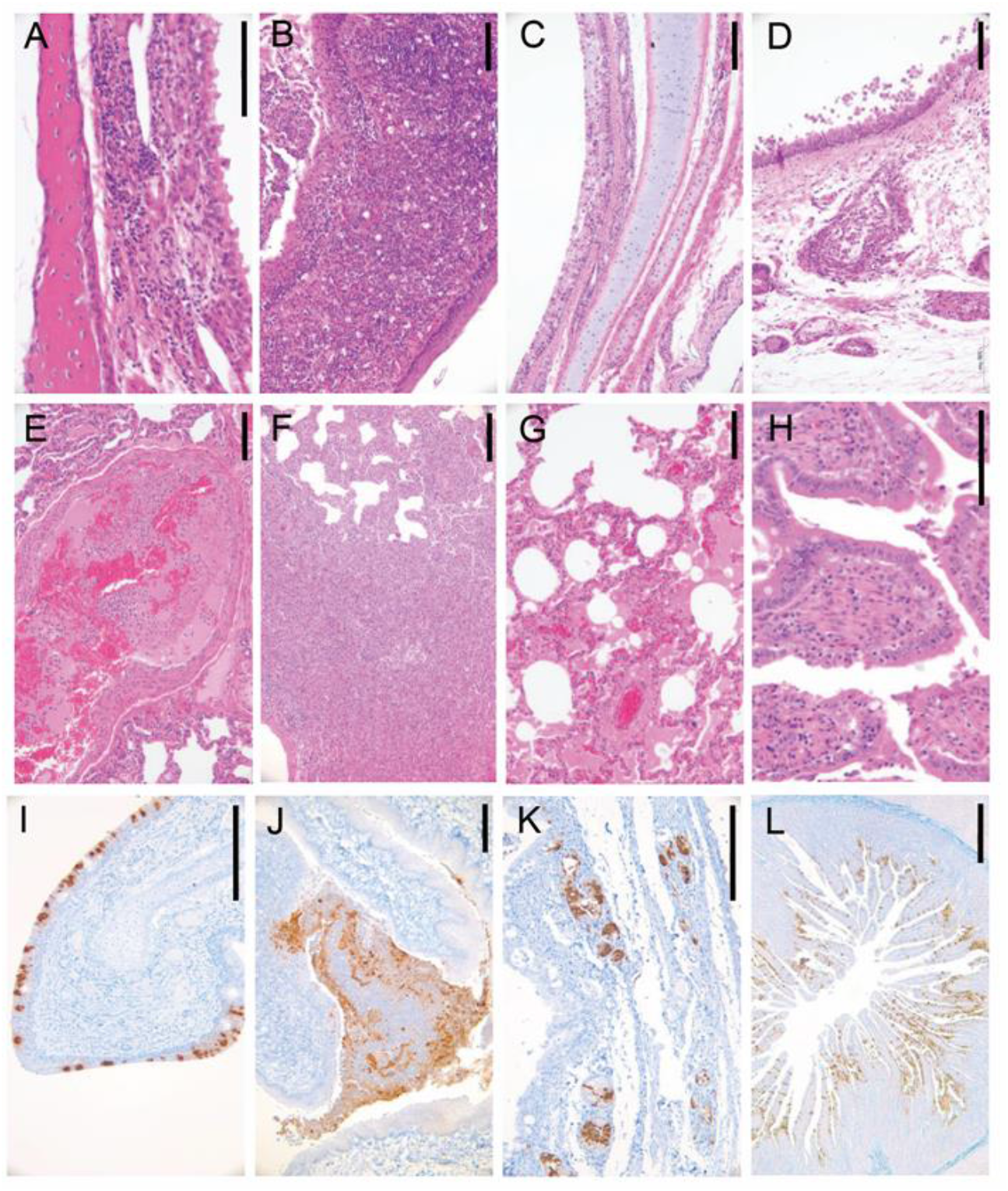
Histological lesions in juvenile cats caused by SARS-CoV-2 infection. Histopathologic and immunohistochemical studies were performed on samples from the virus-inoculated juvenile cats that died or euthanized on day 3 p.i. (**A**) Nasal respiratory mucosa, abnormal arrangement of the epithelium with loss of cilia and lymphocytic infiltration into the lamina propria. (**B**) Tonsil, epithelial degeneration and necrosis with numerous neutrophils and cellular debris on the surface of the epithelium. (**C**) Trachea, degeneration and necrosis of epithelial cells accompanied by coverage with cellular debris and mucous material on the surface of the mucosa. (**D**) Tracheae, epithelial necrosis and lymphocyte infiltration in the submucosal glands. (**E**) Lung, inflammatory cell aggregation and fibrin formation within a blood vessel. (**F**) Lung, infiltration of a large number of macrophages and lymphocytes into the alveolar spaces and interalveolar septa. (**G**) Lung, intra-alveolar edema and congestion in the interalveolar septa. (**H**) Small intestine, viral antigen in the epithelial cells of the mucosa. (**I**) Nasal turbinate, viral antigen in the epithelial cells of the respiratory mucosa. (**J**) Tonsils, viral antigen in the cellular debris and some of the epithelial cells. (**K**) Trachea, viral antigen in the serous cells of the submucosal glands. (**L**) Small intestine, viral antigen in the epithelial cells of the mucosa. Scale bar in A, H=100μm, in B-G, I-K=200μm, in L=500μm.

**Figure S6.**
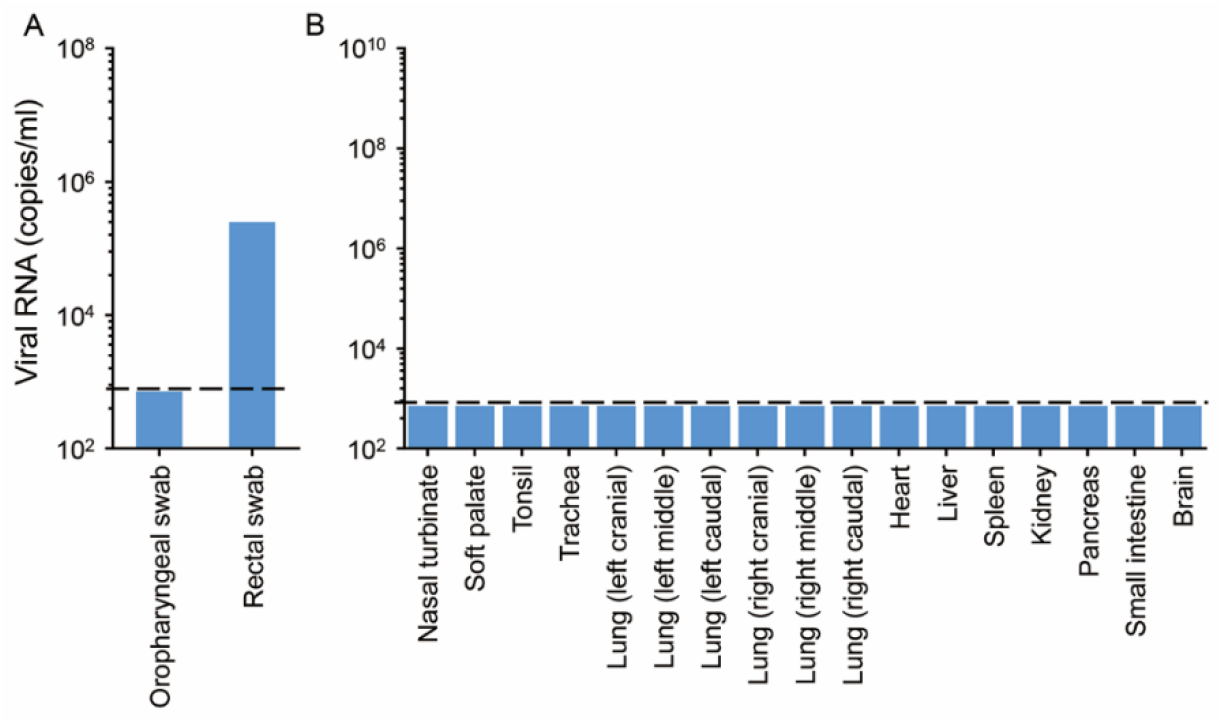
Viral RNA detection in swabs and organs of a dog infected with CTan virus. **(A)** Viral RNA in swabs collected on day 2 p.i. **(B)** Viral RNA in organs or tissues of a dog that was euthanized on day 4 p.i.

**Figure S7.**
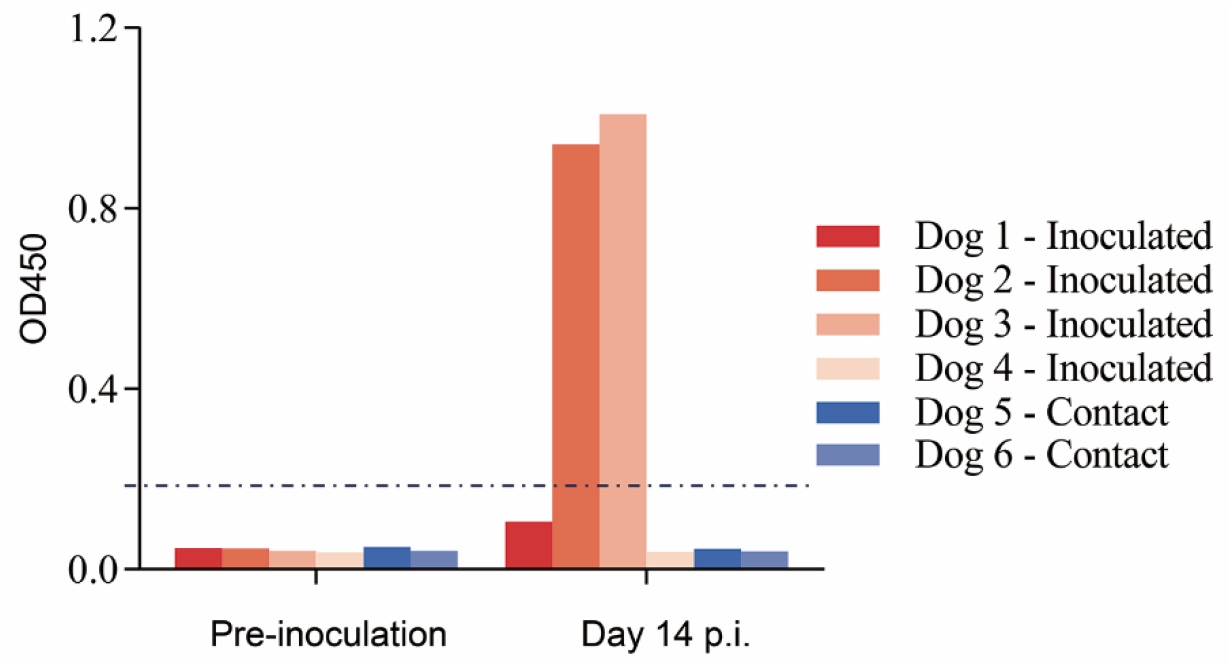
Seroconversion of CTan-H-inoculated and contact dogs. Six-week-old beagles were inoculated with 10^5^ PFU of Ctan-H. Uninfected animals (contact) were housed in the same room with their infected counterparts to monitor the transmission of CTan-H. Sera were collected from animals before inoculation and on day 14 p.i., and antibodies against SARS-CoV-2 were detected by using a Double Antigen Sandwich ELISA kit (ProtTech, Luoyang, China). Optical density values of greater than 0.2 were considered positive for seroconversion according to the manufacturer’s instructions.

